# Brain Activation-based Filtering: a Scholarly Journal Recommendation Model for Human Neuroimaging Studies

**DOI:** 10.1101/2020.10.06.327684

**Authors:** Junsol Kim

## Abstract

The development of noninvasive neuroimaging techniques such as functional magnetic resonance imaging was followed by a large volume of human neuroimaging studies of mental processes, mechanisms, and diseases. Due to the high volume of studies and the large number of journals, it is increasingly challenging for neuroscientists to review existing scholarly journals and find the most suitable journal to publish their studies. Therefore, this paper proposes a scholarly journal recommendation model for human neuroimaging studies called brain activation-based filtering (BAF). Based on the collective matrix factorization technique, BAF recommends journals relevant to the activated brain regions that are described in a given neuroimaging study. For instance, if ‘social brain’ regions such as the dorsomedial prefrontal cortex, precuneus, and temporoparietal junction are activated in a study, BAF recommends relevant social neuroscience journals (e.g., *Social Cognitive and Affective Neuroscience*). Five-fold cross-validation shows that BAF predicts journals with a reliable area under the curve score of 0.855. Furthermore, an interactive Google Colab notebook is offered to recommend relevant journals for a novel human neuroimaging study (https://github.com/JunsolKim/brain-activation-based-filtering).

## 1. Introduction

The development of non-invasive neuroimaging techniques such as functional magnetic resonance imaging (fMRI) was followed by a large volume of human neuroimaging studies of mental processes, mechanisms, and diseases (1). Every year, more than 200 journals publish approximately 6,000 human neuroimaging studies on diverse mental processes, mechanisms, and diseases (2). Therefore, it is increasingly challenging for neuroscientists to review the large body of existing journals and find the most suitable journal to submit their studies to. To reduce the cost of finding a relevant journal, this paper proposes a scholarly journal recommendation model for human neuroimaging studies called brain activation-based filtering (BAF).

The model proposed in this study, brain activation-based filtering (BAF), recommends journals relevant to the activated brain regions that are described in a study. To date, previous studies utilized two strategies to recommend scholarly journals that are based on (1) journals relevant to the authors’ previous publication history (i.e., collaborative filtering) (3–10) and (2) journals relevant to the topics appearing in the text (e.g., keywords, abstract, full paper, and bibliography), like the Elsevier Journal Finder (i.e., content-based filtering) (8,11–16). However, there are several limitations to these strategies. The first strategy is limited by the cold-start issue, which means that journals cannot be recommended to researchers without previous publications (17). The second approach relies on the content of the study. Therefore, journals cannot be recommended if the paper is not written yet. In addition, texts are naturally unstructured, so it is challenging to correctly identify the topics in the text and find relevant journals. To address the issues of these strategies, BAF recommends journals relevant to the activated brain regions that are described in the study. This information is available to researchers without a publication history before the paper is actually written (i.e., right after statistical analyses) and is based on a standardized format (e.g., Montreal Neurological Institute (MNI) template) (19).

To recommend journals based on activated brain regions, BAF uses a machine learning technique called collective matrix factorization (CMF), which is a popular technique for document recommendation models (e.g., news article recommendation models) (20–23). From the Neuroquery database (2), BAF collects the associations between studies and activated brain regions (i.e., which voxel is activated in each study) and the associations between studies and journals (i.e., which journal publishes each study). BAF then constructs a study-voxel association matrix and a study-journal association matrix, which are subsequently decomposed simultaneously via CMF. By doing so, studies and journals are embedded in a common low-dimensional latent topic-space based on activated brain regions. Based on this, journals relevant to each study can be recommended (20–23). For example, if ‘social brain’ regions such as the dorsomedial prefrontal cortex, precuneus, and temporoparietal junction are activated in a study (Van Overwalle, 2009), the study would be related to social neuroscience journals such as *Social Cognitive and Affective Neuroscience* in the latent topic-space. Therefore, social neuroscience journals are recommended for the target study. Such an approach of embedding neuroimaging studies in the latent topic-space based on activated brain regions has long been applied in the previous meta-analysis literature (19,24,25).

This paper consists of four sections. The second section describes the BAF method. The third section presents the experimental results to verify the predictive performance of BAF. The fourth section summarizes the strength and limitation of BAF.

## 2. Materials and Methods

In this section, the BAF model is described. BAF recommends scholarly journals relevant to brain regions activated in a study. To implement BAF, first, two matrices are constructed based on the Neuroquery database (2): (1) the study-voxel association matrix *Y* that indicates which voxels are activated in each study and (2) the study-journal association matrix *Z*, which indicates which journal publishes each study. To evaluate the model, we constructed a training dataset *Z*_*train*_ where 80 % of randomly selected elements from *Z* are included and a test dataset, *Z*_*test*_ where the remaining 20 % elements were included.

Subsequently, as shown in Fig. 1, BAF factorizes the study-voxel matrix *Y* into two latent topic spaces having common dimensions: a study-topic space *S* and voxel-topic space *V*. By doing so, the study-topic space *S* represents the topics of a study and the extent to which a study belongs to these topics (26,27). Second, BAF factorizes the study-journal matrix *Z*_*train*_ into a study-topic space *S* and journal-topic space *J*. Note that the identical study-topic space *S* is used following the CMF technique (20). Finally, BAF trains each latent topic space *S*, *V*, and *J*. Following the training process, the model is validated by examining whether the journal of each study in *Z*_*test*_ is accurately predicted by the study-topic space *S* and the journal-topic space *J*.

**Fig 1.**
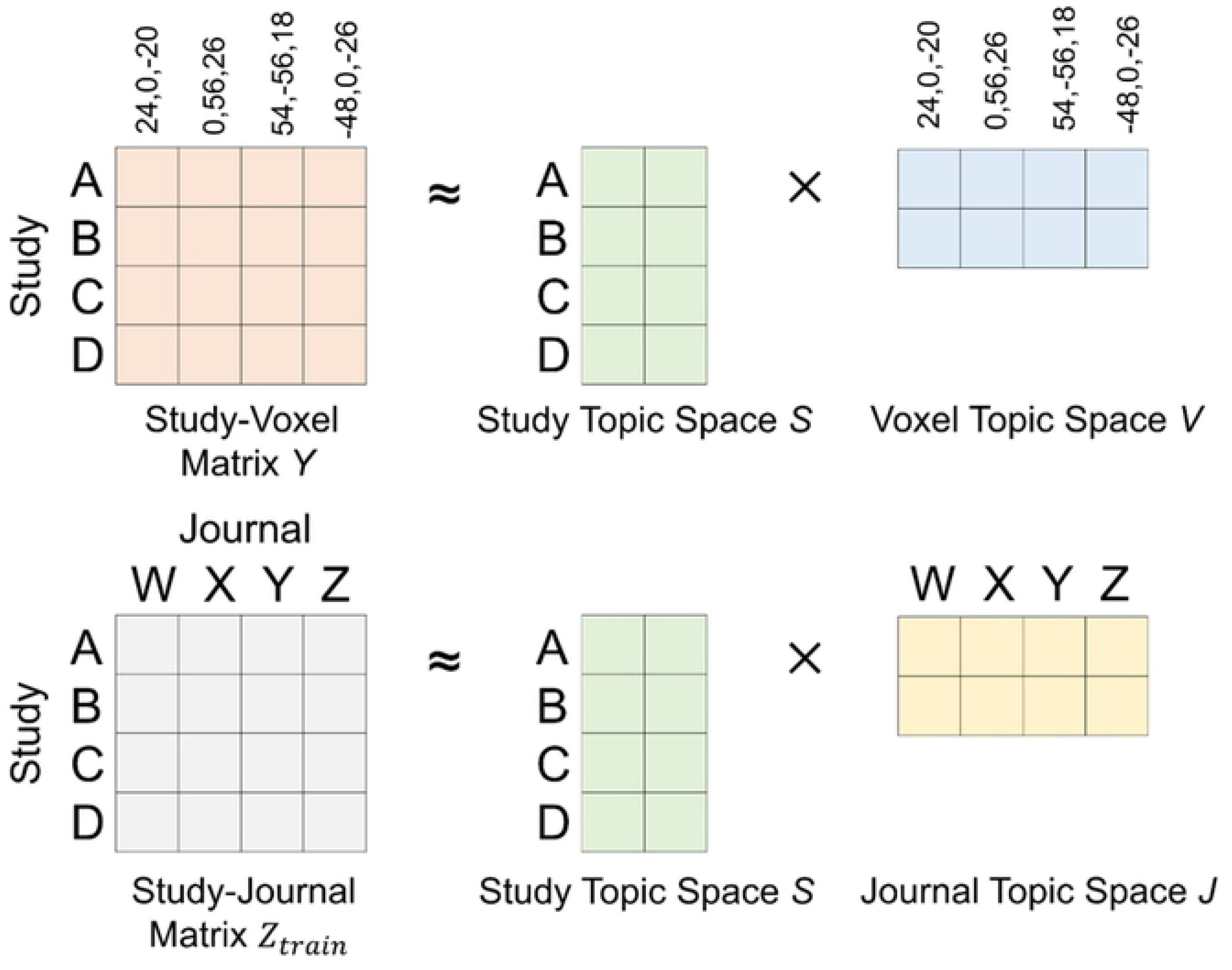
Collective matrix factorization technique.

### 2.1. Data

BAF uses the Neuroquery database (2) which is a recently published public database of activated brain regions in human neuroimaging studies. The data are collected by selecting papers where peak voxel coordinates are extractable and which were published in neuroimaging journals or with query strings such as ‘fMRI’ (2). Journals of the studies in the Neuroquery database were identified using Entrez Programming Utilities at the National Center for Biotechnology Information (28). After discarding papers in journals with fewer than three papers in the database, there were 461,135 peak voxel coordinates activated in 13,173 studies, which were published in 191 journals. Based on these data, (1) the study-voxel association matrix *Y* and (2) study-journal association matrix *Z* were constructed. *N*_*s*_ denotes the number of studies (i.e., 13,173) and *N*_*v*_ denotes the number of possible voxels in the MNI space (i.e., 91 × 109 × 91; MNI-152 2 × 2 × 2 template).

First, we define the study-voxel matrix, 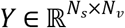 representing an activated voxel *v* in study *s* as follows:

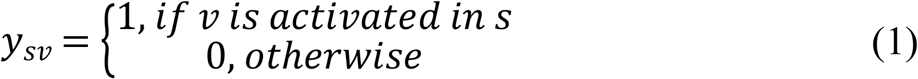

Here, *y*_*sv*_ =1 indicates that voxel *v* is activated in study *s*. We assume that voxel *v* is activated in study *s* if *v* is within 8 mm from the peak voxel coordinates of study *s,* which is in the range of smoothing kernels that are typically used for fMRI meta-analyses (1,2,29–31). Note the non-negativity of *y*_*sv*_. A value of 0 for *y*_*sv*_ does not necessarily mean that voxel *v* is *not* activated in study *s*. Some voxels that are not in the proximity of peak voxel coordinates may have been activated in study *s* (32). Matrix factorization techniques, including the CMF technique, have long been used to process such non-negative data (33).

Second, let *N*_*j*_ denote the number of journals (i.e., 191). We define the study-journal matrix 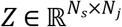 representing that journal *j* publishes study *s* as

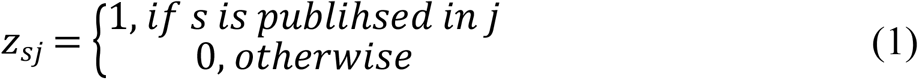

Here, a value of 1 for *Z*_*sj*_ indicates that study *s* is published in journal *j*. However, a value of 0 for *Z*_*sj*_ does not necessarily mean that journal *j* is completely irrelevant to study *s* (i.e., non-negativity). To evaluate the model, we constructed a training dataset 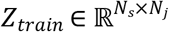 where 80 % of randomly selected elements from *Z* were included and the test dataset, 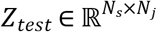 where the remaining 20 % elements were included. We repeatedly split *Z* into *Z*_*train*_ and *Z*_*test*_ for five times so that every element in *Z* is included in *Z*_*test*_ at least once (i.e., five-fold cross-validation).

### 2.2. Collective Matrix Factorization

To obtain the study-topic space *S*, the CMF technique was applied to the BAF model. Matrix factorization techniques have long been used in the field of topic modeling (22,23,26,27) and recommendation models (e.g., Netflix Prize competition) (33,34). Based on the success of matrix factorization techniques in both fields, CMF was developed to build a document recommendation model by jointly implementing topic modeling and recommendation models (20–23).

Based on the CMF technique, BAF maps both studies and activated voxels into two latent topic spaces of dimension *K*, which indicates the number of topics existing in neuroimaging studies and journals (20). We set the value of *K* to 8 to maximize model performance based on an exhaustive search method. Given the original study-voxel matrix *Y*, the goal was to obtain the study-topic space 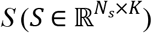 and the voxel-topic space 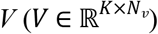. Each row in *S* represents the latent topics of each study and each column in *V* represents the latent topics of each voxel. *S* and *V* were optimized so that the inner product of *S* and *V* was close to the observed entries in the study-voxel matrix *Y.* The journal-topic space *J* can similarly be identified. To identify the journal-topic space *J*, CMF learned the journal-topic space 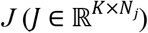 in the way that the inner product of matrices *S* and *J* was close to the observed entries in the study-journal matrix *Z*_*train*_. Note that an identical study-topic space *S* was used (20). By doing so, each column in *J* represents the latent topics of each journal.

### 2.3. Optimization

Three latent topic spaces *S, V, and J* are simultaneously trained to minimize the following cost function (20). Let *S*_*s*_, *V*_*v*_, and *J*_*j*_ denote the latent topic vectors for study *s*, voxel *v*, and journal *j* (i.e., *s*^th^ row of S, *v*^th^ column of V, and *j*^th^ column of J). *y*_*ij*_ and *z*_*ik*_ denote the elements of the study-voxel matrix *Y* and the study-journal matrix, respectively *Z*_*train*_. The cost function of BAF is defined as

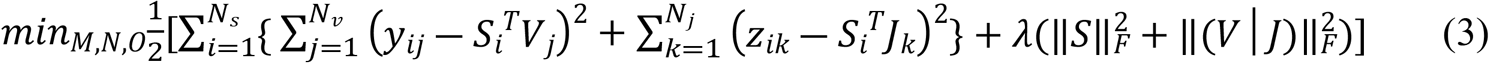

*λ* is a regularization term to avoid over-fitting. (*V*|*J*) is the horizontal concatenation of *V* and *J*. Before training *S, V, and J,* BAF implements random initialization following a Gaussian distribution with mean zero variance. Then, BAF implements the alternate least square technique to train *S, V,* and *J* (33). BAF iteratively alternates between re-computing *S*, *V*, and *J* in the way that each re-computing step lowers the value of the cost function. Specifically, in the first re-computing step, *S* is optimized as follows, while *V* and *J* are fixed.

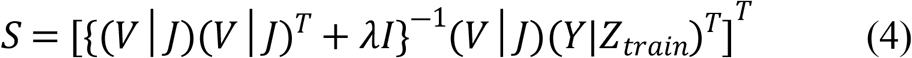

In the second and third re-computing steps, *V* and *J* are optimized as follows, while *M* is fixed. By repeating the first, second, and third steps, the cost function is minimized.

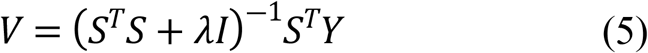

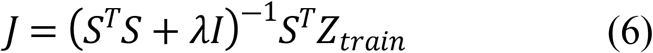

### 2.4. Recommendation of Relevant Journals

After optimizing the study-topic space *S* and journal-topic space *J*, the inner product of the study-topic vector *S*_*s*_ and journal-topic vector *J*_*j*_ can predict the missing values in the original study-journal matrix *Z*_*train*_. If 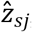, the inner product of *S*_*s*_ and *J*_*j*_, is high, BAF assumes that journal *j* is highly relevant to study *s*. We evaluated the predictive performance of BAF by examining whether 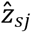 accurately predicts the journals of the studies in the test dataset *Z*_*test*_ (i.e., five-fold cross-validation).

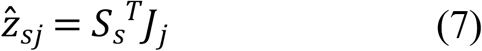

### 2.5. Code Accessibility

An interactive Google Colab notebook to recommend journals for a novel neuroimaging study is freely available online at https://github.com/JunsolKim/brain-activation-based-filtering. In the notebook, activated brain regions in a novel study can be indicated by either peak voxel coordinates (i.e., coordinates of the most activated brain regions) or statistical maps (e.g., T map from the statistical parametric mapping (SPM) toolbox). The experiment code described in this paper is also available online.

## 3. Results

### 3.1. Journal Prediction Performance

To evaluate the performance of BAF, five-fold cross-validation was implemented. As previously mentioned, the study-journal matrix *Z*, which indicates journals that publish the studies, was divided into a training (*Z*_*train*_) and test dataset (*Z*_*test*_). Then, the study-topic space *S* and the journal-topic space *J* were trained using *Y* and *Z*_*train*_ based on the CMF technique. Subsequently, BAF evaluated whether *Z*_*test*_ is accurately predicted by the inner product of *S* and *J* (i.e., 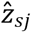). To validate the predictive performance of BAF, we estimated the area under the curve (AUC) score. In addition, we plotted the receiver operating characteristic curve by plotting the false positive rate against the true positive rate based on various thresholds and calculated the AUC score. AUC = 1 means perfect journal prediction by a model and AUC = 0.5 indicates a random selection. The BAF model predicted journals for neuroimaging studies with a reliable AUC score of 0.855.

**Fig. 2.**
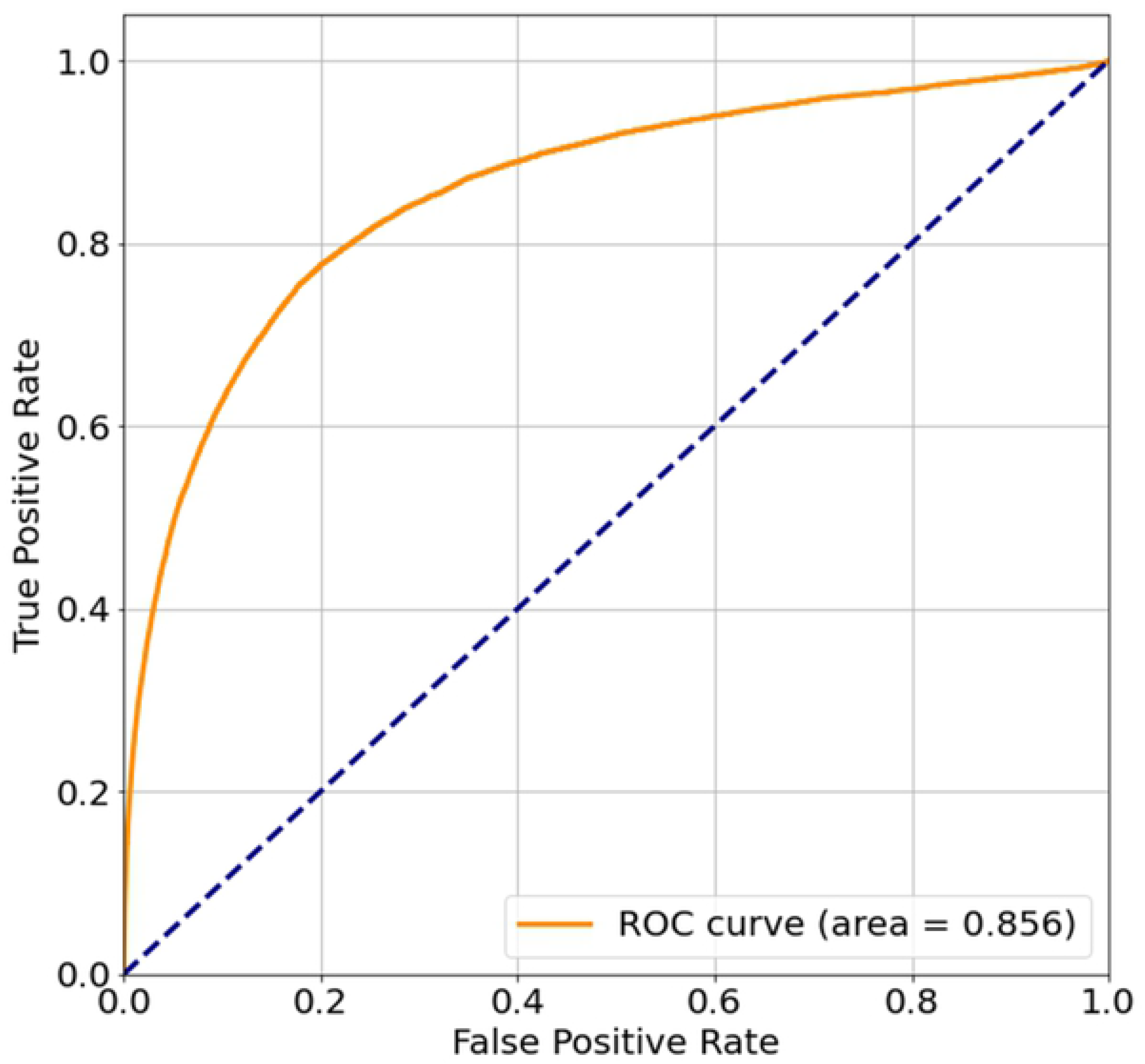
ROC (receiver operating characteristic) curve. Journal prediction performance of brain activation-based filtering (BAF) model is estimated by plotting the ROC curve which plots the false positive rate against the true positive rate based on various thresholds.

### 3.2. Topic Modeling

In the BAF model, all neuroimaging studies and journals are embedded in the common latent topic-space of eight dimensions based on their associated brain activation patterns. By doing so, relevant journals are recommended for neuroimaging studies. Fig. 3 plots the brain regions associated with each latent topic (i.e., each row of voxel topic space *V*), which can be downloaded at https://neurovault.org/collections/8788/. In Table 1, we interpret latent topics by identifying terms associated with brain regions related to each latent topic based on Neurosynth decoder (https://neurosynth.org) (19). Each topic involves the information necessary to identify relevant journals. For instance, Table 1 shows that the first topic is associated with the default mode network, the posterior cingulate cortex (PCC), the medial prefrontal cortex, autobiographical memory, and theory of mind, which are related to social neuroscience. Correspondingly, as shown in Table 2, the journals most associated with the first topic are social neuroscience journals such as *Social Cognitive and Affective Neuroscience* and general human neuroscience journals which publishes a large body of interdisciplinary social neuroscience studies such as *NeuroImage, PLOS One, Neuropsychologia, and Frontiers in Human Neuroscience* (i.e., in the first row of the journal-topic space *J*).

**Fig. 3.**
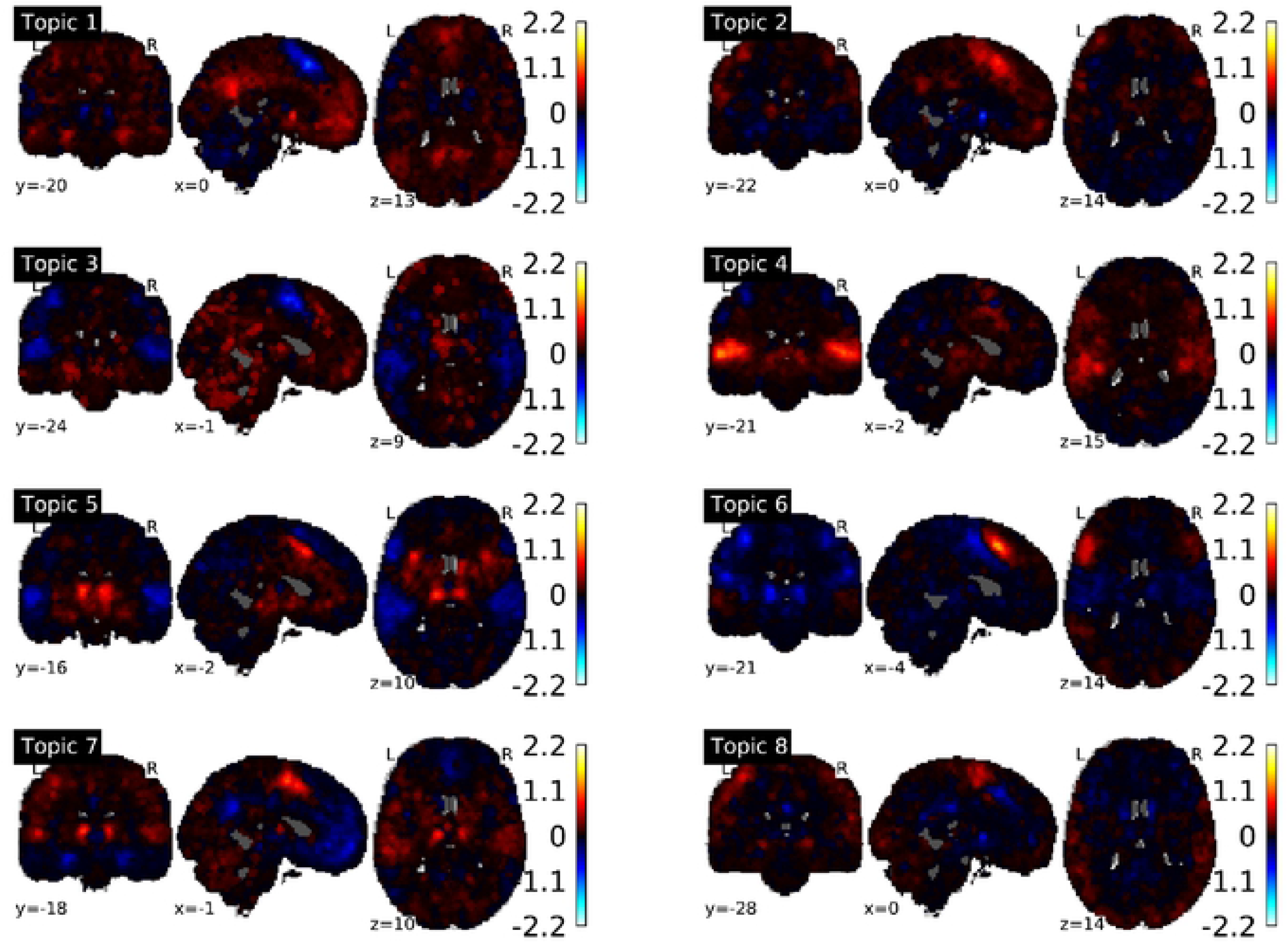
Brain regions associated with latent topics. Each panel presents a map of voxels that are associated with each topic (i.e., voxel latent space *V*).

**Table 1.**
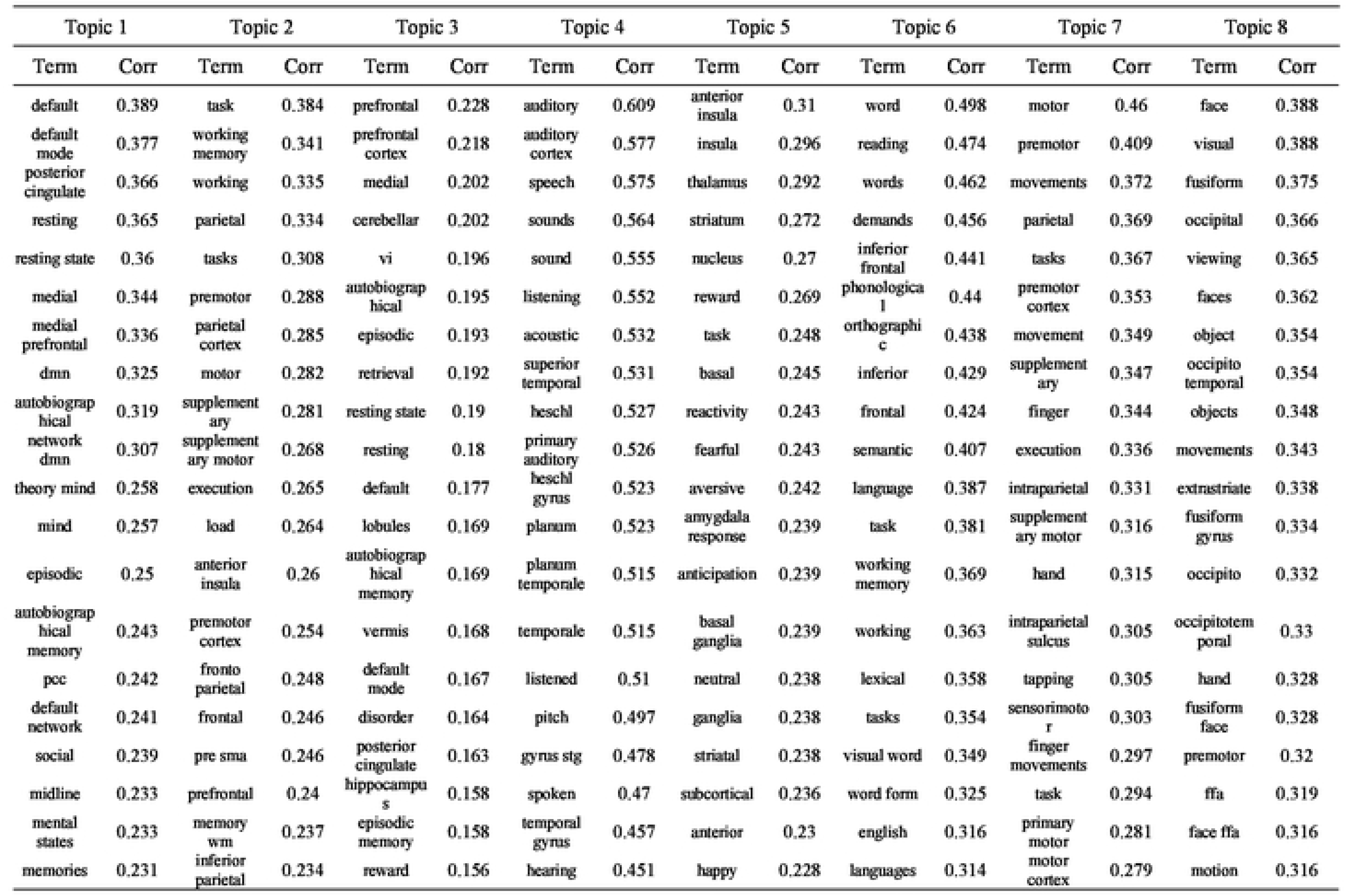
Top twenty keywords associated with latent topics

**Table 2.** Journals associated with latent topics

| Topic | Journals |
| --- | --- |
| Topic 1 | NeuroImage, PloS one, Neuropsychologia, Social cognitive and affective neuroscience, Frontiers in human neuroscience, Brain research, Psychiatry research, Biological psychiatry, NeuroImage. Clinical, Neurobiology of aging |
| Topic 2 | NeuroImage, PloS one, Social cognitive and affective neuroscience, Pain, Neuroscience and biobehavioral reviews, Biological psychiatry, Neuroscience, Brain research, Developmental cognitive neuroscience, Frontiers in systems neuroscience |
| Topic 3 | Social cognitive and affective neuroscience, NeuroImage. Clinical, PloS one, Psychiatry research, Biological psychiatry, Journal of affective disorders, Neurobiology of aging, Neuropsychologia, Frontiers in behavioral neuroscience, Behavioural brain research |
| Topic 4 | NeuroImage, Brain and language, The Journal of neuroscience, Psychiatry research, Biological psychiatry, Social cognitive and affective neuroscience, Pain, Frontiers in psychology, Journal of cognitive neuroscience, Cortex |
| Topic 5 | NeuroImage, Social cognitive and affective neuroscience, Biological psychiatry, Psychiatry research, Pain, PloS one, Biological psychology, Neuron, Human brain mapping, Behavioural brain research |
| Topic 6 | Neuropsychologia, Brain and language, Cerebral cortex, Journal of cognitive neuroscience, Cortex, Brain research, Frontiers in human neuroscience, Frontiers in psychology, Brain research. Cognitive brain research, Human brain mapping |
| Topic 7 | NeuroImage, Brain and language, Neuropsychologia, Magnetic resonance imaging, Brain research, Frontiers in psychology, Brain research. Cognitive brain research, Human brain mapping, Neuroscience, Brain and cognition |
| Topic 8 | NeuroImage, Frontiers in human neuroscience, PloS one, Social cognitive and affective neuroscience, Neuropsychologia, Brain research, Frontiers in psychology, Brain and language, Cerebral cortex, Cortex |

Based on topic mapping, the relations between journals can also be revealed. As shown in the dendrogram in Fig. 4, we characterized the relationships between the largest journals in the Neuroquery database by clustering the journals based on latent topics using hierarchical clustering (single-linkage clustering method). While the figure reveals the similarity between journals based on the traditional division of subjects (e.g., psychiatry), it also reveals novel and interesting similarities between journals. For instance, as expected, psychiatric journals such as *Biological Psychiatry*, *Psychiatry Research*, and *Journal of Affective Disorders* are clustered. However, surprisingly, *Social Cognitive and Affective Neuroscience* is clustered together with such psychiatric journals. In BAF models, the identification of unexpected relations between heterogeneous journals may allow researchers to search for the most suitable journals while avoiding the bias caused by researchers’ previous knowledge about the traditional division of subjects. Furthermore, the dendrogram reveals the hierarchical structure of human neuroscience journals from journals related to specific topics (e.g., *Biological Psychiatry, Social Cognitive and Affective Neuroscience*) to the journals related to general topics (e.g., *Human Brain Mapping*).

**Fig. 4.**
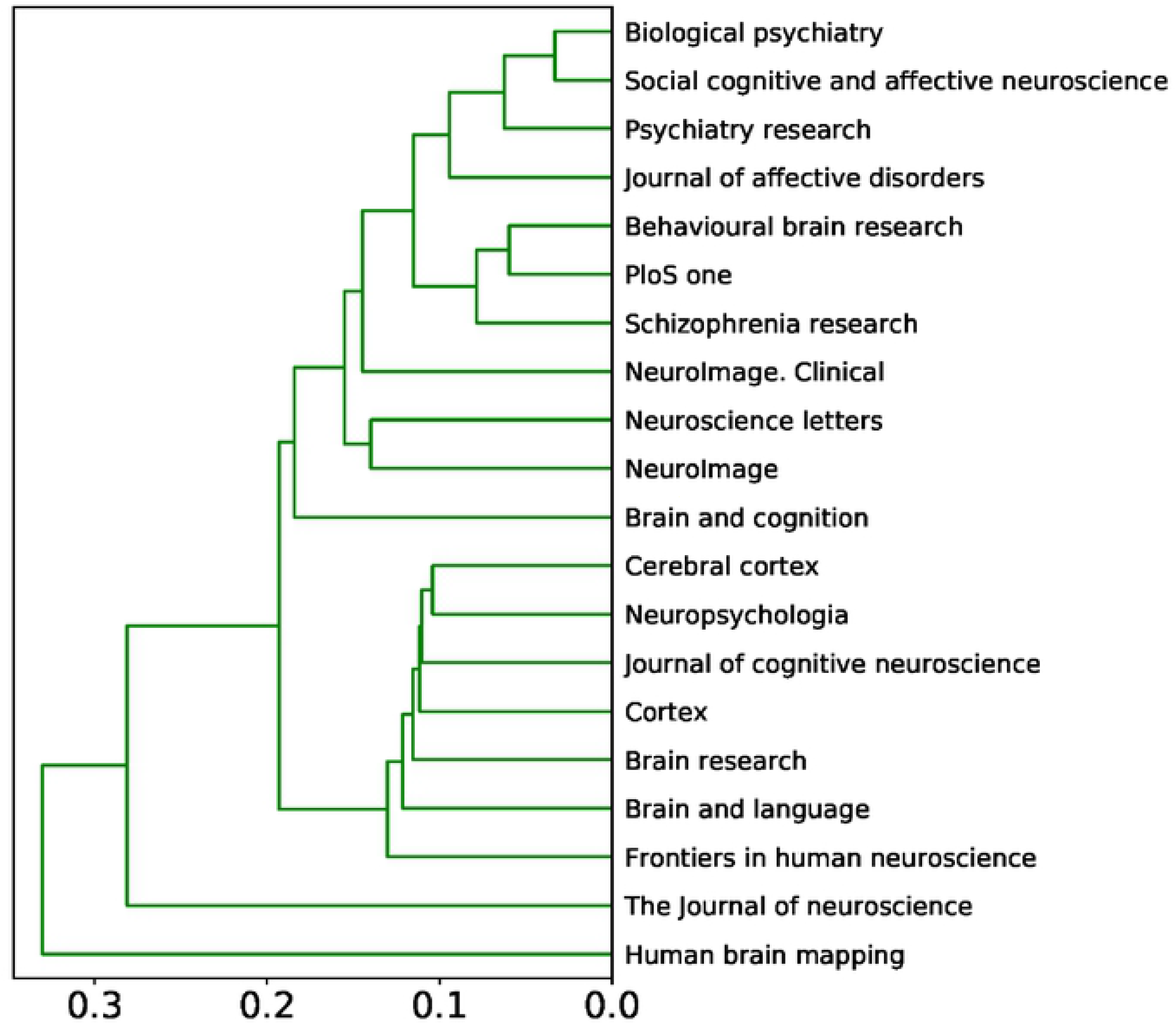
A clustering dendrogram showing the relationships between journals based on their cosine similarity in the latent topic-space. Cosine similarity was used as the distance metric for clustering. Hierarchical clustering was performed using a single-linkage clustering method.

## 4. Discussion

In this study, we propose a novel scholarly journal recommendation model for neuroimaging studies called BAF, which is a method to recommend journals relevant to activated brain regions in neuroimaging studies. The study shows that BAF achieves reliable journal prediction performance (AUC = 0.855) based on five-fold cross-validation. By embedding neuroimaging studies and journals in the common latent space having eight dimensions, BAF could reliably recommend relevant journals to neuroimaging studies, which range from general neuroscience journals such as *NeuroImage* to specialized journals such as *Social Cognitive and Affective Neuroscience*.

Previous neuroinformatics studies have already shown that the topics of neuroimaging studies, such as mental processes, mechanisms, and diseases, can effectively be identified based on activated brain regions. For instance, Nielsen and colleagues identified the topics of 121 neuroimaging studies based on peak voxel coordinates, using a matrix factorization model, which is well-aligned with general neuroscientific knowledge (24). Similarly, Poldrack et al. identified the topics of 5,809 neuroimaging studies based on peak voxel coordinates using a latent Dirichlet allocation model (25). More recently, Rubin and colleagues showed that the topics of neuroimaging studies can be decoded by statistical maps indicating activated brain regions based on neuroimaging databases such as Neurosynth (19). The BAF model extends the previous studies by showing that activated brain regions can be used not only for topic identification and meta-analyses but also for journal recommendations for human neuroimaging studies. In addition, by using activated brain regions, the BAF model overcomes the disadvantages of previous scholarly journal recommendation models, i.e. the author cold start issue, the incapability to recommend journals before writing papers, and the unstructured nature of text data.

However, this study has several limitations. First, as the first study that uses activated brain regions to recommend journals, this study aims to show that activated brain regions alone can be a significant predictor of relevant journals. Although brain activation alone shows reliable performance, the model would be further improved by incorporating other features used in previous scholarly journal recommendation models such as publication history, title, abstract, bibliography, and the full paper. Second, some recently published studies use deep learning-based techniques such as convolutional neural networks (CNNs) when predicting diseases based on brain regions (e.g., brain tumor classification) and achieve excellent performance (35). Future studies may use CNN instead of CMF in the BAF model to test if CNNs improve predictability (36), although matrix factorization-based techniques such as CMF may be superior to CNNs in terms of model efficiency.

## Acknowledgements

This study was supported by the ICONS (Institute of Convergence Science), Yonsei University.

